# The *VEGF* G-quadruplex forming promoter is repaired via long-patch BER

**DOI:** 10.1101/2023.06.25.546439

**Authors:** Adil Hussen, Haley L. Kravitz, Bret D. Freudenthal, Amy M. Whitaker

## Abstract

In response to oxidative damage, base excision repair (BER) enzymes perturb the structural equilibrium of the *VEGF* promoter between B-form and G4 DNA conformations, resulting in epigenetic-like modifications of gene expression. However, the mechanistic details remain enigmatic, including the activity and coordination of BER enzymes on the damaged G4 promoter. To address this, we investigated the ability of each BER factor to conduct its repair activity on *VEGF* promoter G4 DNA substrates by employing pre-steady-state kinetics assays and *in vitro* coupled BER assays. OGG1 was able to initiate BER on double-stranded *VEGF* promoter G4 DNA substrates. Moreover, pre-steady-state kinetics revealed that compared to B-form DNA, APE1 repair activity on the G4 was decreased ∼2-fold and is the result of slower product release as opposed to inefficient strand cleavage. Interestingly, Pol β performs multiple insertions on G4 substates via strand displacement DNA synthesis in contrast to a single insertion on B-form DNA. The multiple insertions inhibit ligation of the Pol β products, and hence BER is not completed on the *VEGF* G4 promoter substrates through canonical short-patch BER. Instead, repair requires the long-patch BER flap-endonuclease activity of FEN1 in response to the multiple insertions by Pol β prior to ligation. Because the BER proteins and their repair activities are a key part of the *VEGF* transcriptional enhancement in response to oxidative DNA damage of the G4 *VEGF* promoter, the new insights reported here on BER activity in the context of this promoter are relevant toward understanding the mechanism of transcriptional regulation.

## Introduction

G-quadruplexes (G4s) have been implicated in the regulation of key cellular processes including DNA replication, recombination, and gene expression (Wang & Vasquez, 2023). At the heart of the G4 structure is a quartet of Hoogsteen-bonded guanines, Figure 1A. The G-quartet helps stabilize the overall G4 structure, which is centrally coordinated by monovalent cations, typically K^+^ ions, and π-π base stacking interactions between tetrads (Figure 1B). Early models assumed a putative quadruplex sequence (PQS) for forming a G4 motif of G_X_-N_1–7_-G_X_-N_1–7_-G_X_- N_1–7_-G_X_, where X ranges from 3 to 6 and N corresponds to any nucleotide (A, G, T or C) forming intermediate loops. Bioinformatics analyses following the completion of the Human Genome Project in the early 2000s predicted that there are hundreds of thousands of PQSs in the human genome (Huppert & Balasubramanian, 2005; Todd et al., 2005). Subsequently, high-throughput sequencing of DNA G-quadruplex structures (G4-seq) in the presence of G4-stabilizing ligand pyridostatin (PDS) identified more than twice as many G4 forming sequences with longer loop lengths and bulges resulting from breaks in G-stretches (Chambers et al., 2015; Marsico et al., 2019). More recently, however, the mapping of G4s by chromatin immunoprecipitation and sequencing (ChIP-seq) using the anti-G4 antibody, BG4, only revealed ∼1000 - 10,000 G4s (Hansel-Hertsch et al., 2016; Hansel-Hertsch et al., 2018). Therefore, the number of G4s that form in chromatin account for only ∼1% of the PQSs identified by bioinformatics and studies using G4 stabilizing ligands. These data strongly support the notion that not all sequences with G4 potential form bonified G4 structures in a particular cellular context. Instead, G4s seem to fold under certain conditions such as upon specific stress stimuli, during certain stages of the cell cycle, in a cell type-specific manner, and/or upon interaction with proteins that bind and stabilize or resolve them (Biffi et al., 2013; Lago et al., 2021; Shu et al., 2022; Varshney et al., 2020). One specific example related to this study is the folding of G4s in the presence of oxidative DNA damage and repair (Cogoi et al., 2018; Fleming et al., 2017; Fleming et al., 2019; Pastukh et al., 2015; Roychoudhury et al., 2020; Wang et al., 2022).

**Figure 1:**
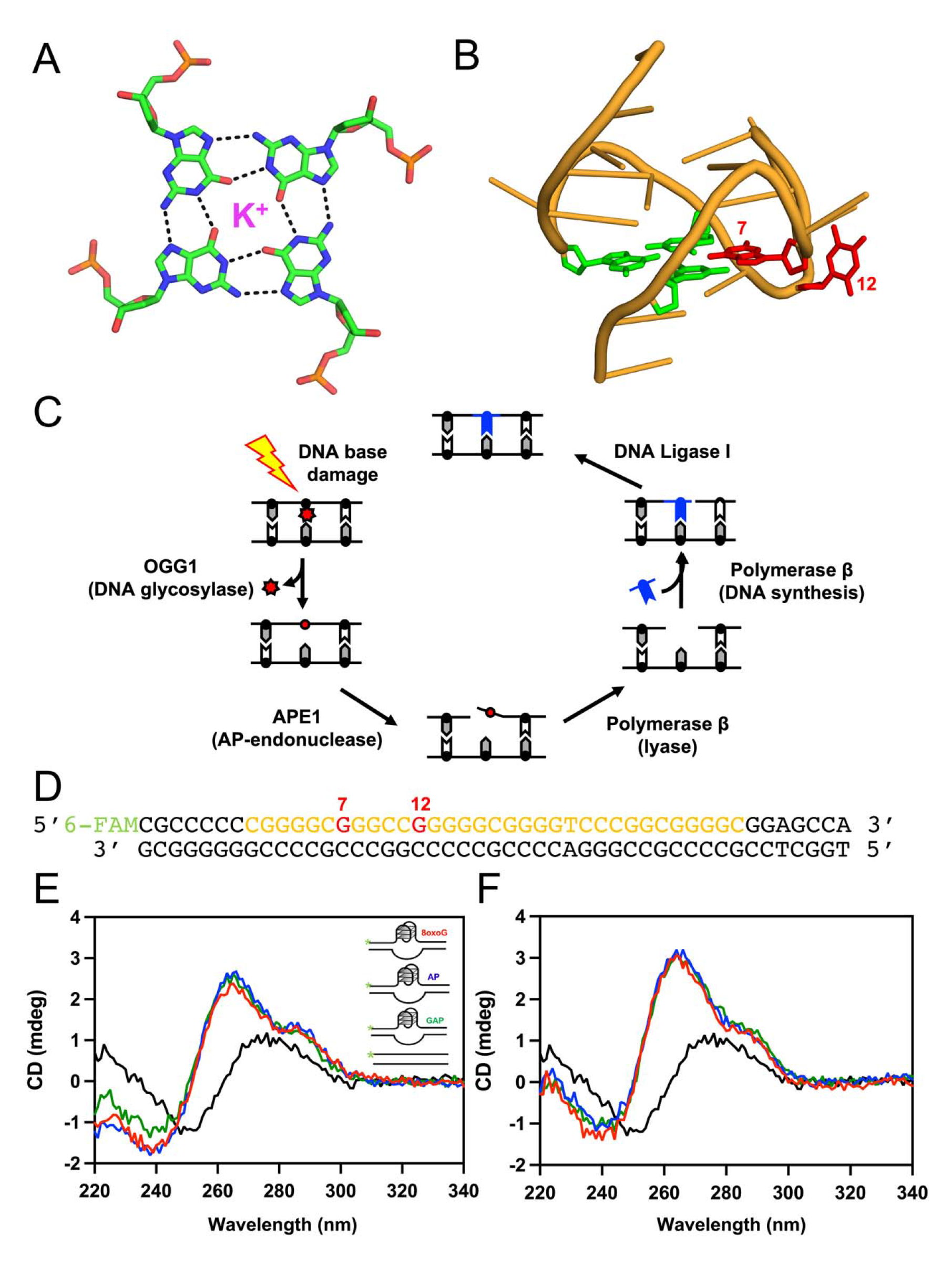
BER intermediates resulting from oxidation of the *VEGF* promoter G4. **(A)** Structure of a G-quartet, showing hydrogen bonds and potassium ion. **(B)** NMR structure of the primary 4 G-runs of the *VEGF* promoter G4 with a G-quartet shown in green and the sites of oxidation damage studied here colored in red (PDB ID 2M27). **(C)** A schematic demonstrating the main enzymatic activities of the BER pathway that remove and replace the oxidative lesion 8oxoG (red star). **(D)** The double-stranded *VEGF* promoter G4 oligonucleotide substrate used for this study. The portion of the sequence in orange represents the 5 G-run *VEGF* PQS, the red Gs indicate damage positions 7 and 12, and the 5’-6FAM fluorescent label used to visualize substrate and product during the *in vitro* BER assays is shown in green. **(E)** CD spectra of the double-stranded *VEGF* promoter oligonucleotide substrates representing BER intermediates with damage at position 7, with red being the substate with 8oxoG, blue a THF, and green a 1-nt gap. Black is a B-form DNA control with a THF. **(F)** CD spectra of the double-stranded VEGF promoter oligonucleotide substrates representing BER intermediates with damage at position 12, with red being the substate with 8oxoG, blue a THF, and green a 1-nt gap. Black is a B-form DNA control with a THF.

Of the 4 DNA nucleobases, G is the most electron rich, and therefore, the most prone to oxidation (Seidel et al., 1996). In fact, Gs at the 5’ end of a G run, such as those in PQS sequences, are even more susceptible to oxidative DNA damage, due to their already low oxidation potential being even further reduced by the additional Gs on their 3’ side (Delaney & Barton, 2003). The primary oxidation product of G is the DNA lesion 8-oxo-7,8-dihydroguanine, 8oxoG (De Bont & van Larebeke, 2004; Fraga et al., 1990). 8oxoG is both a prevalent and mutagenic DNA lesion with potential to initiate and promote disease, including cancer. Fortunately, the well-characterized base excision repair (BER) pathway can defend the cell against 8oxoG, Figure 1C. In the simplest model, BER is initiated when the lesion is removed by a damage-specific DNA glycosylase (e.g., 8-oxoguanine DNA glycosylase or OGG1) to yield an abasic (AP) site. The AP site is then recognized by apurinic/apyrimidinic endonuclease (APE1), which nicks the phosphodiester backbone 5’ of the AP site. APE1 cleavage is followed by the dual AP-lyase and gap filling synthesis activities of DNA polymerase β (Pol β) and the repair cycle is completed via ligation of the remaining DNA nick. While the BER pathway has been well defined in the context of duplex, B-form DNA, it remains unclear how oxidative lesions are repaired in the context of alterative DNA structures, including oxidation-sensitive G4 DNA.

Spatially, G4s are not randomly distributed throughout the genome. Instead, they are enriched at gene promoters, particularly in oncogenes (Eddy & Maizels, 2008; Halder et al., 2009; Huppert & Balasubramanian, 2007), supporting their role in regulating gene-expression and their potential to be targeted for therapeutic intervention. One such example is the *VEGF* promoter, which is transcriptionally activated in response to oxidative DNA damage (Clark et al., 2012; Fleming et al., 2017; Pastukh et al., 2015). This was first shown by qRT-PCR and ChiP analyses, which demonstrated an association between the onset of hypoxia-induced VEGF mRNA expression and the rapid but transient accumulation of 8oxoG in the *VEGF* PQS (Clark et al., 2012; Pastukh et al., 2015). Importantly, ChIP analyses demonstrated that this induced formation of 8oxoG was accompanied by recruitment of BER enzymes OGG1 and APE1 as well as strand-breaks. Later, it was demonstrated in cell-based assays using a plasmid-based luciferase reporter system that 8oxoG in the *VEGF* promoter PQS results in a 2 to 3-fold activation of *VEGF* transcription depending on the specific site of guanine oxidation (Fleming et al., 2017). This site-specific 8oxoG dependent transcriptional activation depended on both the activity of BER glycosylase OGG1 to convert the 8oxoG into an AP site and on the presence of APE1. Of note, with the plasmid-based system, when the repair activity of APE1 is inhibited gene enhancement is further increased, indicating the repair activity of APE1 is not required for gene activation. Moreover, biochemical characterization of APE1 binding and catalytic activity on single-stranded *VEGF* promoter G4 oligonucleotide substrates indicate that APE1 binds the *VEGF* G4 DNA motif with high affinity, however its repair activity is hindered (Fleming et al., 2021). This has been proposed to allow for APE1 to instead recruit gene-activating transcription factors (Fleming & Burrows, 2021). In this model, BER continues to completion through the activities of Pol β and a DNA ligase only after the G-quadruplex is resolved. However, another report shows the opposite effect (i.e., decreased *VEGF* mRNA levels) in response to APE1 strand-cleavage inhibition (Li et al., 2019). These conflicting reports on the effect of inhibiting APE1 strand cleavage activity on *VEGF* expression combined with the prior observation that transient strand breaks are associated with hypoxia-induced VEGF mRNA expression (Pastukh et al., 2015) result in an unclear role for BER activity in the mechanism of 8oxoG-induced *VEGF* gene activation.

As described above, a coupling between *VEGF* transcriptional activation and BER occurs in response to oxidative damage in the *VEGF* promoter G4 motif. Therefore, to understand the currently enigmatic mechanism of *VEGF* transcriptional activation in response to BER, one must also understand the mechanism of BER on the *VEGF* promoter G4 DNA motif. Toward this goal, there are several recent studies characterizing the activates of BER proteins OGG1 and APE1 in the context of G4 DNA. However, there are no studies, to our knowledge, looking at the protein binding and repair activities of BER proteins on *VEGF* G4 DNA substates composed of both the G4 containing strand and the opposing (non-G4) strand, which is the biologically relevant substrate. Moreover, it remains completely unclear how later steps of repair (i.e., DNA synthesis and DNA ligation) occur in the context of the *VEGF* G4 promoter sequence. In this study, we set out to elucidate the mechanism of 8oxoG lesion repair in the *VEGF* promoter G4 motif utilizing double-stranded G4-containing oligonucleotide substrates and purified recombinant enzymes representing each of the major steps of the BER pathway.

## Methods

### DNA Sequences

The VEGF G4 dsDNA substates containing 8oxoG or AP site analog THF were made using the following sequences: G4 strand with a 5’-6-FAM (indicated by asterisk), 5’- *CGCCCCCCGGGGC<u>G</u>GGCC<u>G</u>GGGGCGGGGTCCCGGCGGGGCGGAGCCA-3’, where the underlined Gs represent the positions of either 8oxoG or THF; and the opposing strand, 5’- TGGCTCCGCCCCGCCGGGACCCCGCCCCCGGCCCGCCCCGGGGGGCG-3’. The VEGF G4 DNA ssDNA substates containing 8oxoG were made using the following sequence with a 5’-6-FAM (indicated by asterisk), 5’-*CGCCCCCCGGGGC<u>G</u>GGCC<u>G</u>GGGGCGGGGTCCCGGCGGGGCGGAGCCA-3’. The 1-nt gapped VEGF G4 dsDNA substrate with damage at position 7 was made using the following sequences: upstream G4 strand with a 5’-6-FAM (indicated by asterisk), 5’-*CGCCCCCCGGGGC; downstream G4 strand with a 5’-phospate (indicated by p), 5’- pGGCCGGGGGCGGGGTCCCGGCGGGGCGGAGCCA-3’; and the opposing strand, 5’- TGGCTCCGCCCCGCCGGGACCCCGCCCCCGGCCCGCCCCGGGGGGCG-3’. The 1-nt gapped VEGF G4 dsDNA substrate with damage at position 12 was made using the following sequences: upstream G4 strand with a 5’-6-FAM (indicated by asterisk), 5’-*CGCCCCCCGGGGCGGGCC; downstream G4 strand with a 5’-phospate (indicated by p), 5’- pGGGGCGGGGTCCCGGCGGGGCGGAGCCA-3’; and the opposing strand, 5’- TGGCTCCGCCCCGCCGGGACCCCGCCCCCGGCCCGCCCCGGGGGGCG-3’.

The B-form control for the CD experiments and the APE1 pre-steady-state kinetics was made using the following sequences: AP site strand with a 5’-6-FAM (indicated by asterisk), 5’-*CGTTCGCTGATGCGCXCGACGGATCCGCAT-3’, where the X is a THF; and the opposing strand, 5’- ATGCGGATCCGTCGAGCGCATCAGCGAACG-3’. The 1-nt gapped B-form control dsDNA substrate with damage at position 7 was made using the following sequences: upstream G4 strand with a 5’-6-FAM (indicated by asterisk), 5’-*CGCCGCCCTGACC; downstream G4 strand with a 5’-phospate (indicated by p), 5’- pGACCTTGGGCGGAATCCCGGCTGGGCGGAGCCA-3’; and the opposing strand, 5’- TGGCTCCGCCCAGCCGGGATTCCGCCCAAGGTCCGGTCAGGGCGGCG-3’. The 1-nt gapped VEGF G4 dsDNA substrate with damage at position 12 was made using the following sequences: upstream G4 strand with a 5’-6-FAM (indicated by asterisk), 5’-* CGCCCCCCTGGGCGGACC; downstream G4 strand with a 5’-phospate (indicated by p), 5’- p TGGGCGGAATCCCGGCTGGGCGGAGCCA −3’; and the opposing strand, 5’- TGGCTCCGCCCAGCCGGGATTCCGCCCACGGTCCGCCCAGGGGGGCG-3’. The B-form control for the assay looking at Pol β activity on a substrate made by APE1 with damage at position 7 was made using the following sequences: AP site strand with a 5’-6-FAM (indicated by asterisk), 5’- CGCCCCCCTGGGCXGACCTTGGGCGGAATCCCGGCTGGGCGGAGCCA-3’, where the X is a THF; and the opposing strand, 5’- *TGGCTCCGCCCAGCCGGGATTCCGCCCAAGGTCCGCCCAGGGGGGCG-3’. The B-form control for the assay looking at Pol β activity on a substrate made by APE1 with damage at position 12 was made using the following sequences: AP site strand with a 5’-6-FAM (indicated by asterisk), 5’- *CGCCCCCCTGGGCGGACCXTGGGCGGAATCCCGGCTGGGCGGAGCCA-3’, where the X is a THF; and the opposing strand, 5’- TGGCTCCGCCCAGCCGGGATTCCGCCCACGGTCCGCCCAGGGGGGCG-3’. All sequences containing 8oxoG lesions were purchased from Midland Scientific, and all other sequences were purchased from IDT. The concentration of the individual oligos were determined by absorbance at 260□nm using their extinction coefficients.

### Circular Dichroism

Circular Dichroism spectra were recorded with 400 µL samples at room temperature on an Jasco J-815 spectrophotometer using a 2mm path length quartz cuvette. DNA concentration was 1 µM, and samples were prepared in 50 mM HEPES pH 7.4, 100 mM KCl. 15 scans were accumulated over the wavelength range of 220 – 340 nm at the standard sensitivity, 1 nm data pitch, scanning speed 100 nm min^-1^, 0.5 sec response, and 2 nm bandwidth. Buffers alone were scanned, and these spectra were subtracted from the average scans for each sample. To form the G4 and B-substrates, oligos were annealed at 40 µM in a 1:1 (O8, AP) or 1:1:1 (GAP) ratio in 100 mM KCl, 50 mM HEPES pH 7.4 by heating to 95°C and cooling slowly in a thermocycler. G4-substrates additionally had 100 mM KCl and 15% PEG 200 added to their annealing buffer to facilitate G4 formation (Broxson et al., 2014).

### Expression and Purification of Recombinant BER Proteins

Full-length, wild-type OGG1 was expressed with a GST tag from a pGEX-6P1clone purchased from GenScript. The plasmid was transformed into T7 Express plysS Competent E. coli cells (NEB). Cells were grown in 2xYT at 37°C until OD600 was 0.6. The cells were then induced by adding isopropyl β-D-thiogalactopyranoside (IPTG) to a concentration of 1 mM. The temperature was then reduced to 18°C and cells were allowed to continue growing while shaking overnight before being harvested. After harvesting, cells were lysed at 4°C by sonication in 50 mM HEPES, pH 7.4, 150 mM NaCl, 1 mM EDTA, 1 mM DTT, and a protease inhibitor cocktail (1µg/mL leupeptin, 1 mM benzamidine, 1 mM AEBSF, 1 µg/mL pepstatin A). Lysate was clarified from cell debris by centrifugation for 1 hour at 24,424*g*. The pellet was discarded, and the resulting supernatant was passed over a GSTrap column (GE Health Sciences) equilibrated with lysis buffer (50 mM HEPES, pH 7.4, 150 mM NaCl, 1mM DTT, 1 mM EDTA). OGG1 was eluted from the column with GST elution buffer (50 mM HEPES pH 7.4, 150 mM NaCl, 1 mM DTT, 1 mM EDTA, 50 mM glutathione). Fractions containing OGG1 were then buffer exchanged into low salt buffer (50 mM HEPES pH 7.5, 50 mM NaCl, 1 mM DTT, 1 mM EDTA) and loaded onto a Resource S cation exchange column (Cytiva) equilibrated in the same buffer. OGG1 was eluted with high salt buffer (50 mM HEPED pH 7.5, 1 M NaCl, 1 mM DTT, 1 mM EDTA) and fractions containing OGG1 were buffer exchanged into storage buffer (50 mM HEPES pH 7.4, 150 mM NaCl, 1 mM DTT, 1 mM EDTA). The GST tag was then cleaved using PreScission Protease at 4°C for 6 hours, and cleavage was confirmed via SDS-PAGE gel. After removing the GST tag, OGG1 was concentrated and loaded on a S200 Gel Filtration column (Cytiva). Pure fractions were pooled together, and the final concentration was determined by a NanoDrop One UV-Vis Spectrophotometer (Thermo-Scientific) at an absorbance of 280 nm. Protein was aliquoted and stored at −80°C in a buffer of 50 mM HEPES pH 7.4, 150 mM NaCl, 1 mM DTT, and 1 mM EDTA.

Full-length, wild-type APE1 was expressed in the absence of any tags from a pet28a codon optimized clone purchased from GenScript. The plasmid was transformed into One Shot BL21(DE3)plysS *E. coli* cells (Invitrogen). Cells were grown in 2xYT at 37°C until OD600 was 0.6. The cells were then induced by adding IPTG to a concentration of 0.4 mM. The temperature was reduced to 20°C and cells were allowed to continue growing while shaking overnight before being harvested. After harvesting, cells were lysed at 4°C by sonication in 50 mM HEPES, pH 7.4, 50 mM NaCl, 1 mM EDTA, 1 mM DTT, and a protease inhibitor cocktail. Lysate was clarified from cell debris by centrifugation for 1 hour at 24,424*g*. The pellet was discarded, and the resulting supernatant was passed over a HiTrap Heparin HP column (Cytiva) equilibrated with lysis buffer (50 mM HEPES, pH 7.4, 50 mM NaCl). APE1 was eluted from the column using a linear gradient of NaCl up to 600 mM. APE1 was then buffer exchanged into 50 mM NaCl and loaded onto a Resource S column (Cytiva) equilibrated in lysis buffer (50 mM HEPES, pH 7.4, 50 mM NaCl, 1 mM DTT, 1 mM EDTA). APE1 was eluted from the column using a linear gradient of NaCl up to 400 mM. APE1 was subsequently loaded onto a HiPrep 16/60 Sephacryl S-200 HR (Cytiva) equilibrated in 50 mM HEPES, pH 7.4, and 150 mM NaCl. SDS-PAGE was used to determine purity of the resulting fractions. Pure fractions were pooled together, and the final concentration was determined by a NanoDrop One UV-Vis Spectrophotometer (Thermo-Scientific) at an absorbance of 280 nm. Protein was aliquoted and stored at −80C in a buffer of 50 mM HEPES, pH 7.4 and 150 mM NaCl.

Human full-length wild-type DNA Pol β was overexpressed from a PET-28a codon optimized clone purchased from GenScript in the BL21-CodonPlus(DE3)-RP *E. coli*. Induction of expression was initiated when the OD600 reached 0.6 at 37C using 0.1 mM IPTG. Temperature was reduced to 18°C and cells were harvested the following day. Purification was carried out as previously described (Beard & Wilson, 1995). In brief, transformed *E. coli* lysate was run over a HiTrap Heparin HP (Cytiva), Resource S (Cytiva) and HiPrep 16/60 Sephacryl S-200 HR (Cytiva) columns. Fractions containing pure Pol β were concentrated and stored at −80°C in 50 mM HEPES pH 7.4, 150 mM NaCl. Pol β was determined to be pure via SDS page and final concentration was determined via A280 using a NanoDrop One UV-Vis Spectrophotometer.

Full-length N-terminal HIS-tagged human wild-type DNA ligase 1 (LIG1) was overexpressed from a pET-15b clone, gifted to us from Dr. Sam Wilson, in BL21- CodonPlus(DE3)-RP *E. coli* cells. Overexpression and purification were carried out as previously described with slight modification (Chen et al., 2006). Induction of expression was initiated when the OD600 reached 0.7 at 37°C using 0.1 mM IPTG. The temperature was reduced to 16°C and cells were harvested 18 hours later. Briefly, purification occurred as follows. Transformed *E. coli* lysate was run over HisTrap HP (Cytiva), HiTrap Heparin HP (Cytiva), Mono Q (Cytiva) and HiPrep 16/60 Sephacryl S-200 HR (Cytiva) columns. Fractions containing pure LIG1 were concentrated and stored at −80°C in 50 mM HEPES pH 7.4, 150 mM NaCl. Ligase 1 was determined to be pure via SDS-PAGE and the final concentration was determined via A_280_ using a NanoDrop One UV-Vis Spectrophotometer.

Purified full-length human wild-type FEN1 protein was provided by Dr. Bill Beard from the lab of Dr. Sam Wilson at the NIEHS. Purification was verified via SDS page, and the final concentration was confirmed via A_280_ using a NanoDrop One UV-Vis Spectrophotometer.

### *In vitro* coupled BER assays

The in vitro BER assays were performed using a benchtop heat block set to 37°C. These assays, which include B-form or G4 substrates (100 nM), were performed at an initial reaction volume of 130 uL in buffer containing 50 mM HEPES pH 7.4, 100 mM KCl, 10 mM MgCl_2_, and 0.1 mg/mL BSA. The specific enzyme(s) used in each assay are indicated on the figures and figure legends, and concentrations were the following: 150 nM OGG1, 25 nM Pol β, 0.5 nM APE1, 100 nM LIG1, and 25 nM FEN1 unless specifically indicated otherwise in the figure legend. Assays containing LIG1 also included 1 mM ATP, and assays containing Pol β included 100 uM dGTP (or other dNTP as indicated). 15 uL aliquots were taken at the indicated times over the course of the 60-minute reaction and immediately quenched by mixing with an equal volume of DNA gel loading buffer (100 mM EDTA, 80% deionized formamide, 0.25 mg ml^−1^ bromophenol blue and 0.25 mg ml^−1^ xylene cyanol). After incubation at 95°C for 5 min, the reaction products were separated by 22% denaturing polyacrylamide gel. A GE Typhoon FLA 9500 imager in fluorescence mode was used for gel scanning and imaging, and the data were analyzed with Image J software (Schneider et al., 2012). To form the G4 and B-form substrates, oligos were annealed at 1 µM in a 1.2:1 (O8, AP) or 1.2:1.2:1 ratio (lower concentration of the 6FAM labeled oligo) in TE by heating to 95°C and cooling slowly in a thermocycler. G4- substrates additionally had 100 mM KCl and 15% PEG 200 added to their annealing buffer to facilitate G4 formation (Broxson et al., 2014).

### Pre-steady-state APE1 AP-endonuclease assays

A rapid quench-flow instrument (KinTek RQF-3) was used for the APE1 AP-endonuclease activity measurements. The reaction buffer was 50□mM HEPES, pH 7.4, 100□mM KCl, 3□mM MgCl_2_, and 0.1□mg□ml^−1^ BSA. Final concentrations were 100□nM DNA substrate and 30□nM APE1 after mixing. At indicated time intervals, aliquots were quenched by mixing with 100□mM EDTA. An equal volume of DNA gel loading buffer (100□mM EDTA, 80% deionized formamide, 0.25□mg□ml^−1^ bromophenol blue and 0.25□mg□ml^−1^ xylene cyanol) was added to the quenched reaction mixture. After incubation at 95□°C for 5□min, the reaction products were separated by 22% denaturing polyacrylamide gel. All time points are the mean of at least three independent experiments. A GE Typhoon FLA 9500 imager in fluorescence mode was used for gel scanning and imaging, and the data were analyzed with Image J software (Schneider et al., 2012). The biphasic time courses were fit to the equation: Product□=□*A*(1−*e*^−kobst^)□+□*v*_ss_*t*, where *A* represents the amplitude of the rising exponential and *k*_obs_ the first order rate constant. The steady-state rate constant (*k*_ss_) is the steady-state velocity (*v*_ss_) *A*^−1^, where *A* represents the fraction of actively bound enzyme. To form the G4 and B-form substrates, oligos were annealed at 1 µM in a 1.2:1 ratio (lower concentration of the 6FAM labeled oligo) in TE by heating to 95°C and cooling slowly in a thermocycler. G4-substrates additionally had 100 mM KCl and 15% PEG 200 added to their annealing buffer to facilitate G4 formation (Broxson et al., 2014).

### Fluorescence polarization binding assays

Fluorescence anisotropy measurements were used to quantify binding of full length wild-type APE1 to FAM-labelled DNA substrates described above. FAM-labelled DNA substrates were mixed with APE1 concentrations ranging from 0 - 512 nM in costar® black polypropylene 96- well round bottom plates. For all titrations, the concentration of the FAM-labelled DNA was 1 nM to stay below the *K*_D_ and maintain sufficient signal to noise. Assembled plates were incubated at room temperature for one hour prior to taking measurements. Fluorescence anisotropy measurements were carried out on a SpectraMax i3x Multi-Mode Microplate reader with a fluorescence polarization fluorescein detection cartridge (FP-FLUO) at 22°C in buffer consisting of 25 mM HEPES pH 7.4, 100 mM KCl, 15 mM EDTA, 10 mM MgCl_2_, and 1 mM DTT. The excitation and emission wavelengths were 485 and 535 nm, respectively. Polarization values were calculated using the following equation:

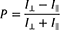

Fluorescence polarization changes were fit by nonlinear least squares (NLS) regression to a one-site specific binding model using Prism (GraphPad). A minimum of four independent measurements were averaged for each APE1 concentration of the titrations to determine *K*_D_ values.

## Results

### VEGF promoter G4 DNA substrates

To determine the activity of the BER enzymes OGG1, APE1, Pol β, and DNA ligase I on DNA containing a damaged *VEGF* promoter G4 motif, we first designed 5’-6FAM labeled double-stranded G4-containing oligonucleotides representing substrates and intermediates of the BER pathway (Figure 1D). Our DNA sequence included the four G-runs primarily involved in G4 formation, plus a fifth G-run 7 nucleotides distant to the sequence in the 3’ direction. This fifth G-run has been hypothesized to act in the role of a “spare tire” for aiding in repair of damaged G4s, including that of the *VEGF* promoter by swapping out a G-run containing a damaged base to place the lesion in a large loop (Fleming et al., 2015). For the oligonucleotide substrates to be able to adopt the G4 fold while also maintaining contact with the opposing C-rich strand we flanked the G4 sequence with 7 additional nucleotides on either end. We choose two damage sites within the *VEGF* G4 sequence that map to core (position 7) and loop (position 12) regions of the G4 based on the solution NMR structure (PDB ID 2M27; Figure 1B) of the G4 *VEGF* promoter sequence (Agrawal et al., 2013). Importantly, positions 7 and 12 are both particularly oxidation prone (Fleming et al., 2015) and exhibit transcriptional activity in response to oxidative damage that is dependent on BER enzymes OGG1 and APE1 (Fleming et al., 2017).

The above-described double-stranded DNA oligonucleotide substates containing G4 sequences can be formed via annealing the G4 forming strand and its C-rich complement in buffer containing 100 mM KCl, 15% PEG 200 to facilitate G4 formation (Broxson et al., 2014). To confirm G4 formation of the substates in the assay buffer, we performed circular dichroism (CD) experiments on each of our double-stranded G4 substates with either an 8oxoG, AP site, or 1-nt gap at position 7 or 12 and looked for spectra characteristics ascribed to G4 DNA. For both position 7, Figure 1E, and position 12, Figure 1F, each substrate presents a λ_max_ of 265 nm with a shoulder at 290 nm and a λ_min_ of 240 nm, characteristic of a hybrid G4s (Lim et al., 2010; Ravichandran et al., 2018). This signature shape is consistent with those seen in the CD spectrum of the single-stranded *VEGF* G4 sequence containing the 5^th^ G-run (Fleming et al., 2015), and importantly confirms that our double-stranded substates are in a G4 conformation under our assay conditions.

### OGG1 activity on the *VEGF* promoter G4 sequence

During the initiation of BER, OGG1 binds 8oxoG and cleaves the N-glycosyl bond that covalently links the nitrogen atom from the amino group of the nucleotide to the anomeric carbon of the ribose sugar, converting 8oxoG into an AP site. As a bifunctional DNA glycosylase, OGG1 also has AP lyase activity, which creates a nick in the DNA by cleaving the phosphodiester bond 3’ of the AP site. Of note, OGG1 is thought to function primarily in a monofunctional mode *in cellulo* leaving strand cleavage to APE1 (Allgayer et al., 2016; Dalhus et al., 2011; Franck et al., 2022; Hill et al., 2001). OGG1 has demonstrated strong substrate specificity, with preference for 8oxoG:C base pairs in double-stranded DNA as opposed to 8oxoG in single-stranded DNA (Franck et al., 2022). In the context of the *VEGF* G4 DNA sequence, weak OGG1 activity has been demonstrated on a single-stranded G4 oligonucleotide containing 5 G-runs but not on a *VEGF* G4 DNA sequence with only 4-runs (Fleming et al., 2015). As this was a single-stranded G4 substate, it was proposed that this activity may result from the formation of a transient hairpin structure between a run of Cs in the sequence and one of the G-runs. Alternatively, the 5^th^ G-run may allow for the adaptation of an alternative G4 structure on which OGG1 is active.

Here, we utilized a *VEGF* G4 DNA sequence containing both the 5^th^ G-run and an opposing strand and compared OGG1 activity to the same *VEGF* G4 DNA sequence in the absence of the opposing strand (Figure 2). In agreement with the previous work, we saw weak OGG1 activity on the substrates lacking the opposing strand with slightly higher activity when the damage is in position 7 (core) compared to position 12 (loop). More surprisingly, we saw greatly enhanced OGG1 activity on the double-stranded G4 DNA, which would mimic the biologically relevant substrate. In fact, at the 60-minute points, a majority of the DNA has been cleaved. DNA cleavage was eliminated when the assay was performed with a catalytically dead OGG1 variant (Supplemental Figure 1), confirming G4 DNA cleavage occurs via the canonical OGG1 active site. All together, these data indicate that either OGG1 can resolve the G4 into canonical B-form DNA prior to performing catalysis and/or the presence of the opposing strand and flanking duplex regions enhance OGG1 catalysis.

**Figure 2:**
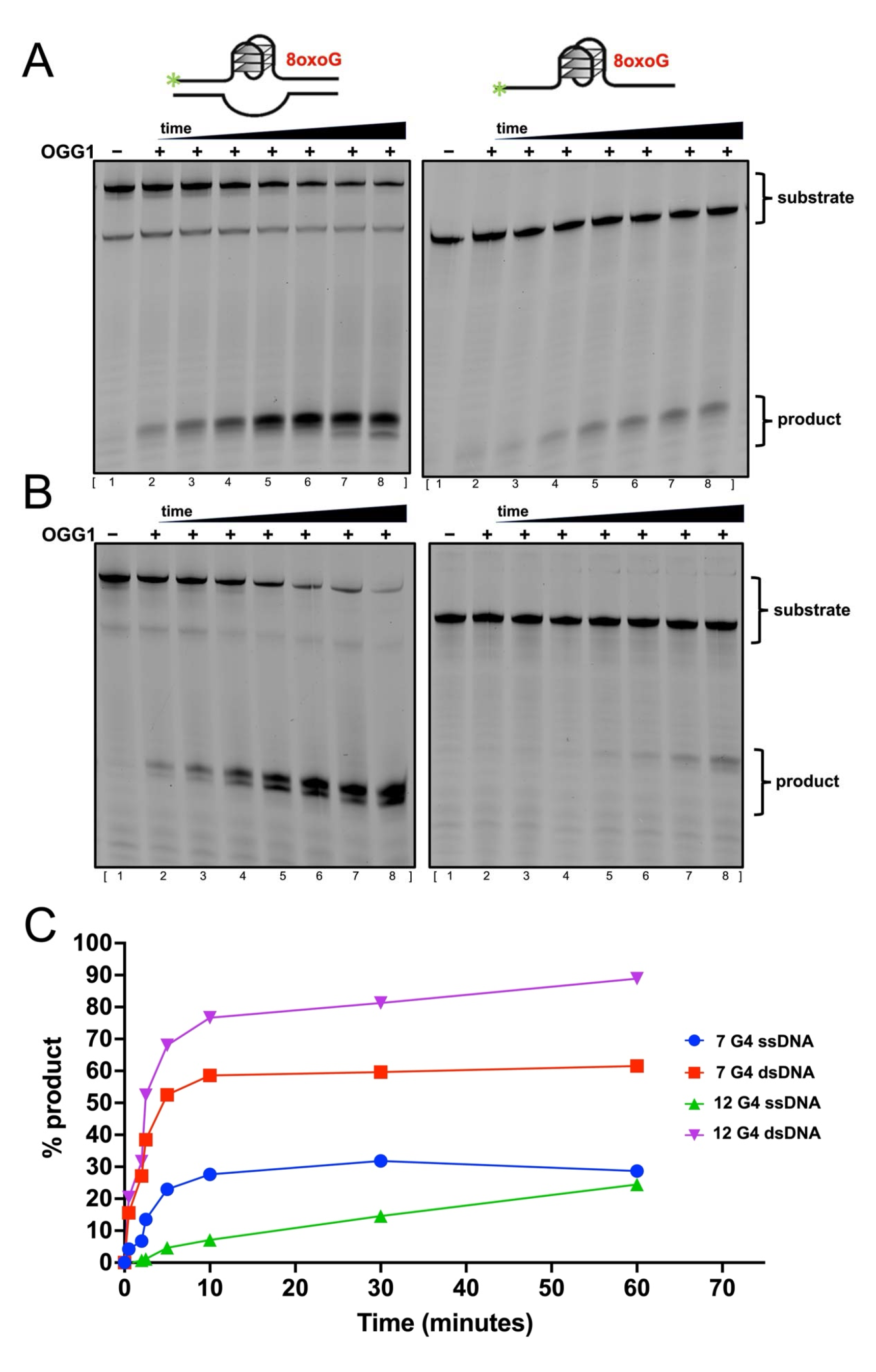
OGG1 cleavage of *VEGF* G4 promoter substates. **(A)** Representative gel images for the OGG1 reaction time courses on the double-stranded (left) and single-stranded (right) *VEGF* G4 substrates with an 8oxoG lesion at position 7. **(B)** Representative gel images for OGG1 reaction time courses on the double-stranded (left) and single-stranded (right) *VEGF* G4 substrates with an 8oxoG lesion at position 12. Lane 1 of each panel is a no enzyme control. Lanes 2 – 8 are the reaction products of OGG1 N-glycosyl bond cleavage followed by phosphodiester backbone incision over a 60-minute time course. The reaction time points corresponding to lanes 2 – 8 are 0.5, 1, 2.5, 5, 10, 30, and 60 minutes. Note that the 8oxoG containing double-stranded substrates often run as two species due to partial re-annealing of the complementary strand to the 6-FAM labeled G4 containing strand during gel loading. **(C)** Quantification of the OGG1 reaction time courses for each substate.

### APE1 activity on the *VEGF* promoter G4 sequence

During BER, APE1 is responsible for the cleavage of the sugar phosphate backbone 5’ to the AP site to produce a single-stranded break with 5’-deoxyribose phosphate and 3’-hydroxyl end groups. This AP-endonuclease activity of APE1 has been previously measured in the context of several G4 sequences, including that of the G4 promoters of *VEGF* (Fleming et al., 2021), *cMYC* (Broxson et al., 2014; Roychoudhury et al., 2020), and *NEIL3* (Fleming et al., 2019; Howpay Manage et al., 2022), as well as telomeric G4 sequences (Burra et al., 2019; Howpay Manage et al., 2023). Overall, the APE1 AP-endonuclease activity or product yield in all these studies is characterized as reduced compared to that seem in B-form duplex DNA substrates. However, the degree of reduced activity is quite variable depending on the assay conditions and unique G4 topography. Of note, apart from one of the studies with the cMYC G4 promoter sequence, all published APE1 binding and activity assays were done in single-stranded G4 DNA as opposed to double-stranded G4 oligonucleotide substates. Therefore, it remains largely unclear how APE1 cleaves double-stranded DNA containing an AP site within a G4 motif with the opposing strand, as would be observed in a cellular context. To address this, we characterized the activity of APE1 on a double-stranded G4 oligonucleotide by pre-steady-state kinetics. We obtained biphasic time courses of product formation for the G4 substrates, demonstrating that as previously observed for B-form DNA substrates, catalysis during the first enzymatic turnover (burst phase) is more rapid than that in the subsequent steady-state phase, where a step after chemistry (likely product release) is rate limiting (Figure 3A). Overall, we observed efficient catalysis of the G4 substrates. Specifically, APE1 cleaves the double-stranded G4 DNA substrates with AP sites at positions 7 and 12 at rates of 39.7 ± 7.0 and 40.9 ± 13 s^-1^, respectively. These rates are about 2-fold faster than the B-form control (k_obs_ = 18.8 ± 4.5 s^-1^). On the other hand, the steady-state rates were 2.3 and 1.6 -fold slower compared to B-form DNA for the G4s substrates with AP sites at positions 7 (k_ss_ = 0.69 ± 0.15 s^-1^) and 12 (k_ss_ = 1.0 ± 0.2 s^-1^), respectively. This indicates that while APE1 cuts the DNA efficiently it does take about twice as long to turnover the reaction in the case of our G4 substrates. In other words, APE1 is slower to release the cleaved G4 product than it is does the B-form product. This slower product release would have implications for the recruitment of transcription factors and time APE1 spends on a G4 motif. These differences do not come from changes in the ability of APE1 to bind the G4 DNA, as the apparent binding affinities of APE1 to the three substates are very similar, Figure 3B.

**Figure 3:**
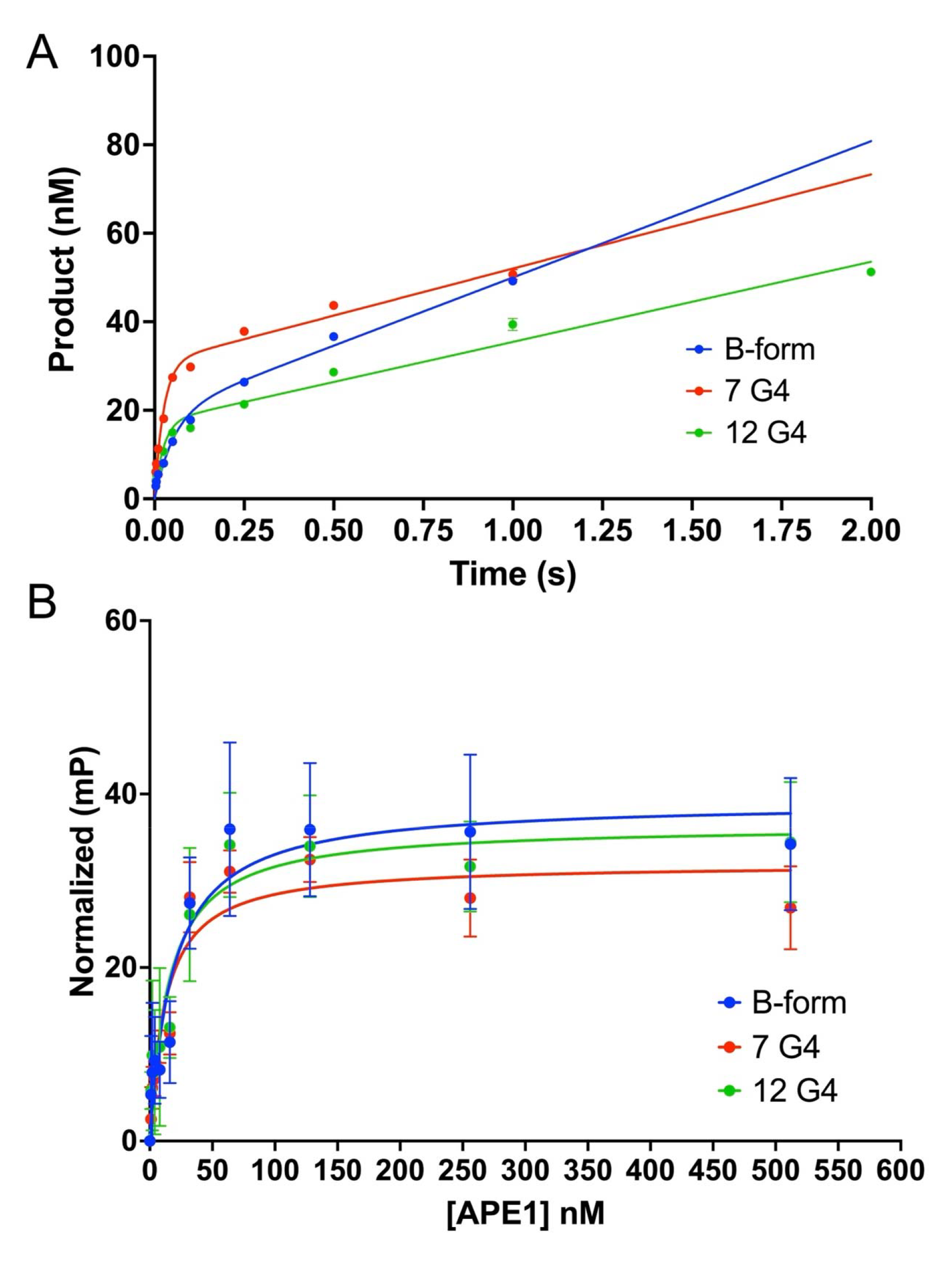
Kinetic and binding characterization of the APE1 AP-endonuclease reaction on *VEGF* G4 promoter substates. **(A)** Pre-steady-state kinetic time courses of product formation are shown for APE1 on an abasic B-form DNA control (blue), a double-stranded *VEGF* promoter G4 substrate with an AP site analog THF at position 7 (red), and a double-stranded *VEGF* promoter G4 substrate with a THF at position 12 (green). **(B)** Binding of APE1 to B-form DNA control substrate with a centrally located AP site analog THF (blue), a double-stranded *VEGF* promoter G4 substrate with an AP site analog THF at position 7 (red), and a double-stranded *VEGF* promoter G4 substrate with a THF at position 12 (green). All error bars represent the standard error of the mean from three replications of the experiment and in some cases the error bars are hidden by the points. Kinetic and binding parameters are shown in Table 1.

**Table 1:**

| | $[APE1]_{\text{active}}$ (nM) | $k_{\text{obs}}$ (s<sup>-1</sup>) | fold change | $k_{\text{ss}}$ (s<sup>-1</sup>) | fold change | $K_D$ (nM) | fold chang |
| --- | --- | --- | --- | --- | --- | --- | --- |
| **B-form** | 19.2 ± 2.1 | 18.8 ± 4.5 | - | 1.6 ± 0.3 | - | 18 ± 5 | - |
| **7 G4** | 30.8 ± 2.0 | 39.7 ± 7.0 | 2.1 ↑ | 0.69 ± 0.15 | 2.3 ↓ | 12 ± 2 | 1.5 ↓ |
| **12 G4** | 17.3 ± 1.7 | 40.9 ± 13 | 2.2 ↑ | 1.0 ± 0.2 | 1.6 ↓ | 15 ± 4 | 1.2 ↓ |

### Pol **β** DNA synthesis on G4 substrates

After AP site cleavage by APE1, Pol β performs both its DNA lyase and DNA synthesis activities. To our knowledge, the mechanism of DNA gap filling by a polymerase within a G4 substrate remains unexplored. Because we were unsure if this activity would be template-dependent, we first tested the ability of Pol β to insert each of the 4 nucleotides (dATP, dTTP, dGTP, dTCP) into double-stranded *VEGF* G4 substrate with a single nucleotide (1-nt) gap at either position 7 or 12, Figure 4. There was very little insertion observed with the pyrimidine nucleotides, moderate insertion of a dATP, and multiple insertions of dGTP. The number of dGTP insertions was dependent on the position of the 1-nt gap in the sequence, with 2 insertions in the case of position 7 (Figure 4A) and 5 in the case of position 12 (Figure 4B). Of note, in the *VEGF* promoter sequence position 7 is followed by 2 Gs and position 12 is followed by 4, Figure 1C. Therefore, the data is consistent with Pol β utilizing the DNA template and continuing to synthesize the DNA until reaching a mismatch, apart from the lack of insertion of a third dGTP in the case of position 7. Since DNA synthesis occurs via extension of the 3’ end of strand upstream the DNA nick, this would indicate that the DNA upstream the site of damage in the 1-nt gapped substrate is able to re-anneal to the DNA template strand.

**Figure 4:**
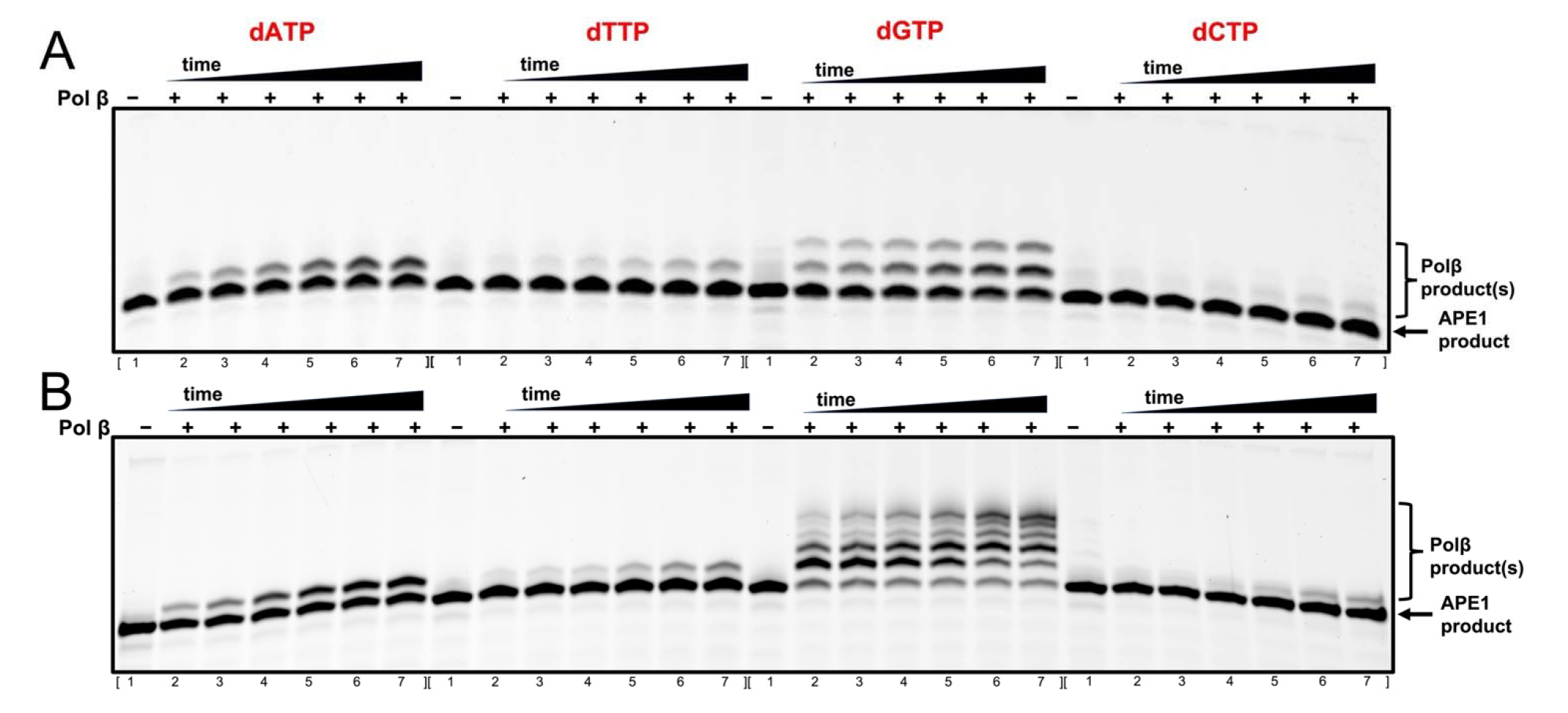
Nucleotide discrimination of the Pol β gap filling activity on *VEGF* G4 promoter substates. **(A)** Representative gel image for a Pol β reaction time course on a double-stranded *VEGF* G4 substrates with a 1-nt gap at position 7. **(B)** Representative gel image for a Pol β reaction time course on a double-stranded *VEGF* G4 substrates with a 1-nt gap at position 12. Lane 1 of each time course is a no enzyme control. Lanes 2 – 7 are the reaction products of Pol B insertion of the indicated dNTP over a 60-minute time course. The reaction time points corresponding to lanes 2 – 7 are 1, 2.5, 5, 10, 30, and 60 minutes.

To further explore Pol β activity on the *VEGF* G4 promoter DNA sequences, we next looked at Pol β insertion of dGTP on substrates generated by the BER pathway enzymes preceding it in the repair pathway, APE1 and OGG1. In these assays the DNA substrates with damage at position 7 (Figure 5A) or position 12 (Figure 5B) were first incubated for 30 minutes with APE1 (AP site substrates) or both APE1 and OGG1 (8oxoG substrates) to generate the APE1 product/Pol β substrate. Next, Pol β was added and dGTP insertion was monitored over time. Unlike the G4 substrates, but as expected, the B-form DNA resulted in a single insertion of dGTP (Figure 5A and 5B, left side). On the other hand, regardless of whether the reaction was started with an AP site by APE1 or an 8oxoG by OGG1/APE1 Pol β DNA synthesis resulted in the same multiple insertion pattern obtained with the 1-nt gapped DNA substrates described above. Of note, APE1 cleavage on the ssDNA G4 substrates, second time course from the left in Figure 5A/B, was poor as had been previously reported (Fleming et al., 2021). APE1 did have some activity on the ssDNA G4 with an AP site at position 7, however the overall yield was low and did not result in any Pol β DNA synthesis products. There was no band present corresponding to an APE1 cleavage product with the corresponding substrate with an AP site at position 12 under the assay conditions. Therefore, we could not initially test Pol β DNA synthesis activity on the ssDNA G4. However, at 100 times the concentration of APE1 some cleaved product was generated on the position 12 ssDNA G4 substate that did not go on to serve as a substrate for Pol β (Supplemental Figure 2).

**Figure 5:**
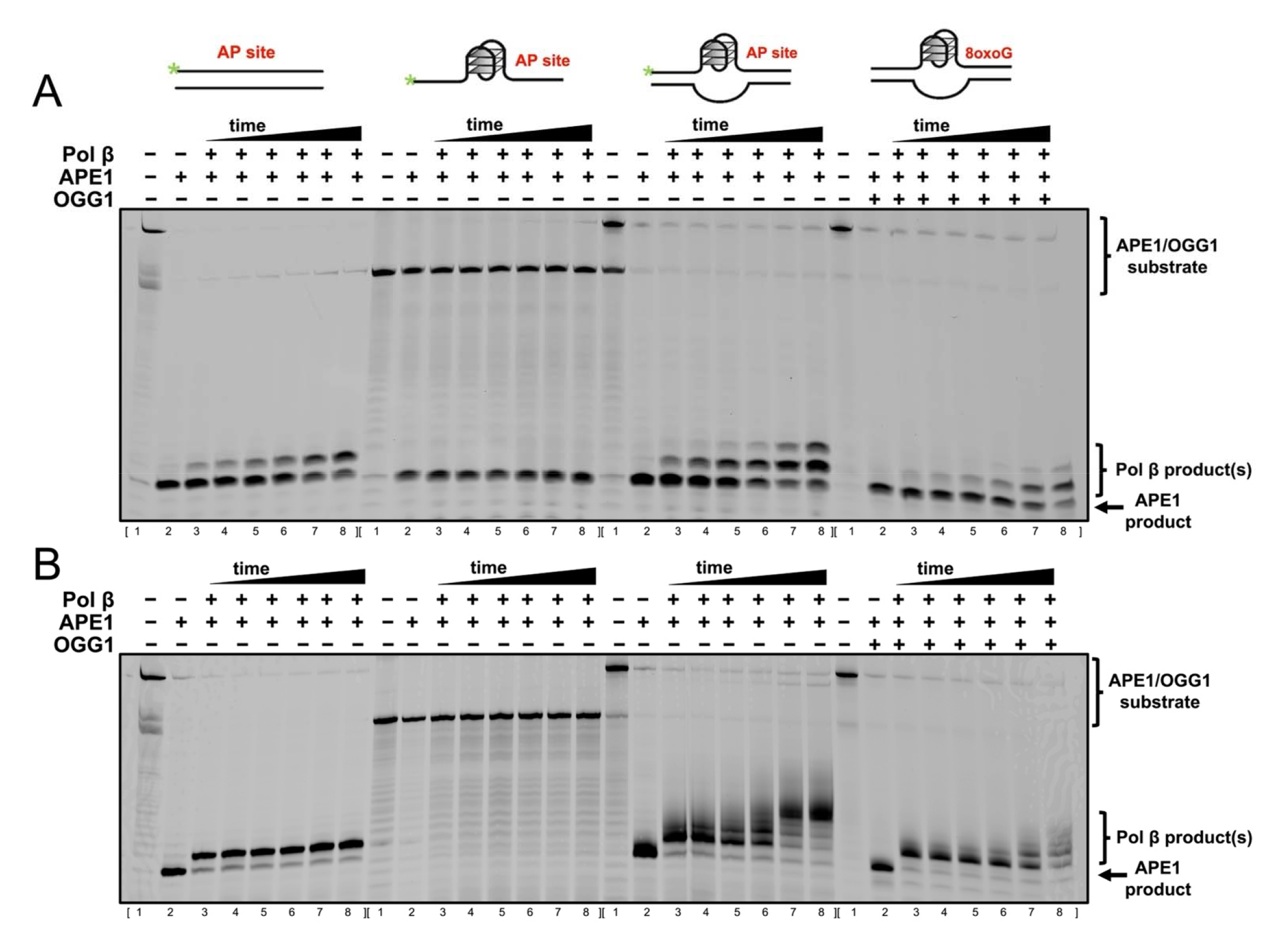
Pol β gap filling activity on *VEGF* G4 promoter substates by a coupled BER assay. **(A)** Representative gel image for a Pol β reaction time course on various *VEGF* substrates with damage at position 7. Substrates from left to right are the following: B-form with AP site analog THF, single-stranded VEGF G4 promoter with a THF, double-stranded VEGF G4 promoter with a THF, and double-stranded VEGF G4 promoter with an 8oxoG. **(B)** Representative gel image for a Pol β reaction time course on various *VEGF* substrates with damage at position 12. Substrates from left to right are the following: B-form with AP site analog THF, single-stranded VEGF G4 promoter with a THF, double-stranded VEGF G4 promoter with a THF, and double-stranded VEGF G4 promoter with an 8oxoG. Substrates with a THF were first incubated with APE1 for 30 minutes to generate a substrate for Pol β, whereas the 8oxoG substrates were first incubated with both APE1 and OGG1. Lane 1 of each time course is a no enzyme control. Lane 2 is after 60-minute incubation with either APE1 or APE1/OGG1. Lanes 3 – 8 are the reaction products of Pol β insertion of dGTP over a 60-minute time course. The reaction time points corresponding to lanes 3 – 8 are 1, 2.5, 5, 10, 30, and 60 minutes. Note that the THF and 8oxoG containing double-stranded substrates often run as two species due to partial re-annealing of the complementary strand to the 6-FAM labeled G4 containing strand during gel loading.

### Ligation depends on FEN1

Following insertion of a single nucleotide, gap DNA synthesis by Pol β in B-form duplex DNA is completed when the resulting nick is ligated. This final step of BER results in the regeneration of non-damaged DNA. To test the ability of DNA ligase 1 to ligate the nicked DNA substrates produced by Pol β during BER of the G4 DNA *VEGF* substrates, we first incubated Pol β with the 1-nt gapped G4 DNA substrate for two hours. This resulted in substrates for DNA ligase in which almost all of the DNA gaps have undergone multiple insertions (Figure 6, lanes with only Pol β). Interestingly, the G4 substrate with a 1-nt gap at position 7 resulted in lower yields of DNA synthesis products than either the G4 with damage at position 12 or the B-form controls. This indicates that this substrate may be more difficult to extend from and/or the strand upstream the DNA nick may have more difficulty re-annealing to the opposing strand. Next, we added DNA ligase 1 and looked for the formation of a ligation product over time. For both sites of damage, the substate with multiple insertions was unable to be ligated (Figure 5A and B, far left time courses), whereas the B-form control resulted in a higher weight ligation product (Figure 5A and B, second time course from the left). Since the substrate sequences contained only a 1-nt gap, the extra insertions likely occur via strand displacement DNA synthesis and would result in a short DNA flap as occurs during long-patch BER, Figure 6A. Therefore, we next tested whether the long-patch associated flap endonuclease, FEN1, could cleave the flap producing a product capable of being ligated by DNA ligase 1, Figure B and C. FEN1 improved ligation in the substrate with damage at position 7 compared to the ligation reaction done in the absence of FEN1 (Figure 6B). However, ligation is still less efficient than in the reactions with B-form DNA. For position 12, Figure 6C, the presence of FEN1 improved the degree of ligation to the same level as the B-form DNA control. In both cases FEN1 appears to have no additional effect on ligation in the B-form substrate (second versus fourth time course in Figure 5A and B).

**Figure 6:**
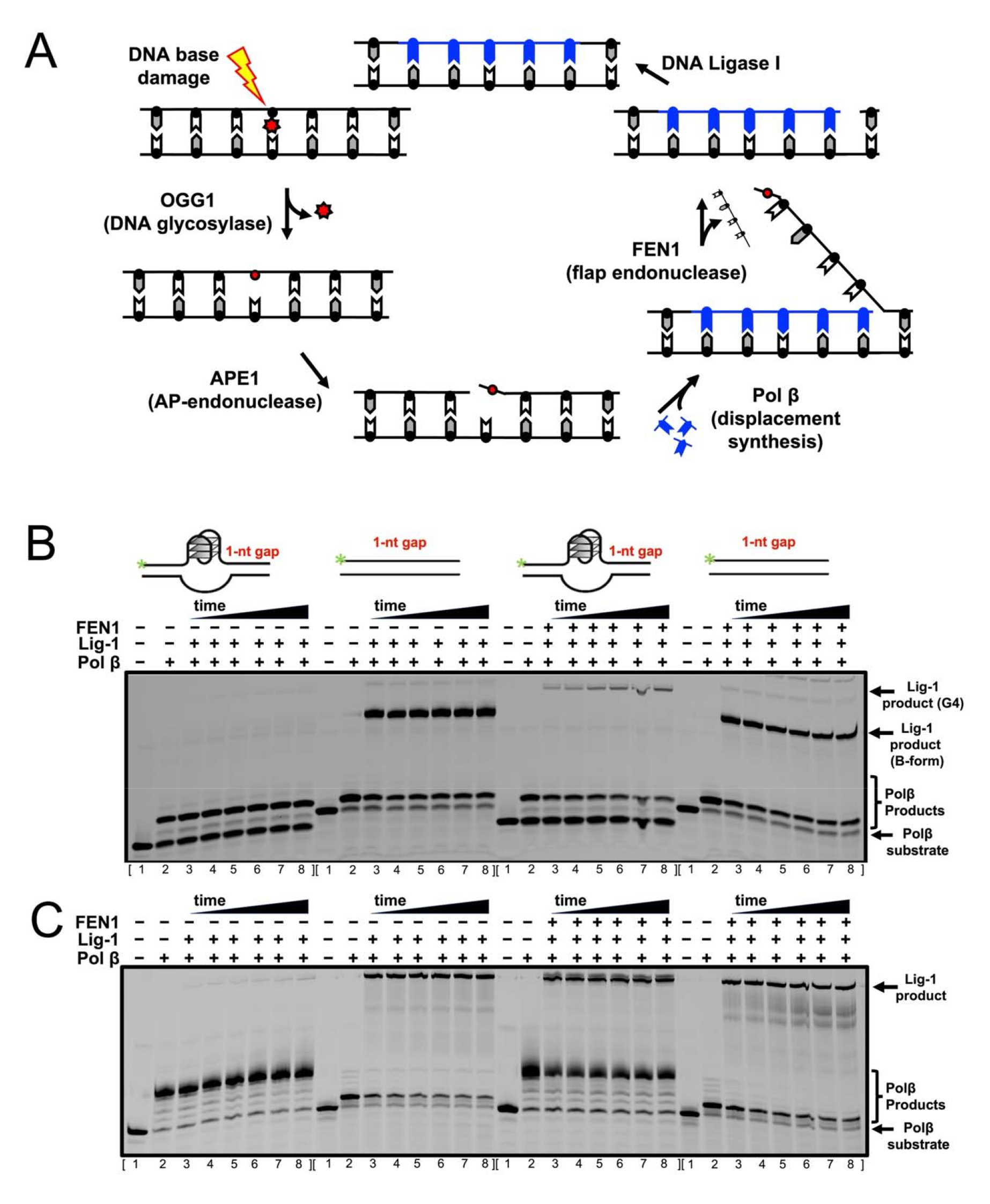
DNA ligase 1 ligation activity on *VEGF* G4 promoter substates by a coupled BER assay. **(A)** Schematic demonstrating long-patch BER. **(B)** Representative gel image for DNA ligase 1 reaction time courses on position 7 1-nt gapped double-stranded *VEGF* promoter substrates in either G4 or B-form. Ligation in the absence of FEN1 is shown in the first two time courses, and ligation in the presence of FEN1 is shown in the second 2 time courses. **(C)** Representative gel image for DNA ligase 1 reaction time courses on position 12 1-nt gapped double-stranded *VEGF* promoter substrates in either G4 or B-form. Ligation in the absence of FEN1 is shown in the first 2 time courses, and ligation in the presence of FEN1 is shown in the second 2 time courses. The 1-nt gapped substrates were first incubated with Pol β for 2 hours to generate a substrate for DNA ligase 1. Lane 1 of each time course is a no enzyme control. Lane 2 is after the 2-hour incubation with Pol β. Lanes 3 – 8 are the reaction products of DNA ligase 1 over a 60-minute time course. The reaction time points corresponding to lanes 3 – 8 are 1, 2.5, 5, 10, 30, and 60 minutes.

## Discussion

The idea that G4s can act as transcriptional regulators originated in 2002 when Hurley *et al*. provided evidence that the G4 in the promoter region of the oncogene c-*MYC* was biologically active by showing c-*MYC* downregulation as a direct consequence of G4 stabilization by the porphyrin-based ligand TMPyP4 (Siddiqui-Jain et al., 2002). This observation suggested that G4s could act as transcriptional repressors, a concept supported in several subsequent studies on G4s formed in other oncogene promoters, such as *c-KIT*, *BCL-2*, *KRAS*, *TERT*, and *VEGF* (Balasubramanian et al., 2011). This general concept of G4s as transcriptional repressors was further strengthened when bioinformatic studies revealed the enrichment of G4s at gene promoters, particularly in oncogenes (Eddy & Maizels, 2008; Halder et al., 2009; Huppert & Balasubramanian, 2007), and G4 structures were consequently promoted as targets of small molecules for therapeutic intervention. Since, this idea of G4s as direct transcriptional repressors has been challenged, as G4 stabilizing ligands were shown to cause gene down-regulation of target genes in response to factors besides target G4 stabilization, including induction of DNA damage at the G4 (Rodriguez et al., 2012) and global G4 stabilization (Boddupally et al., 2012). More recently, new technologies and methods including G4 ChIP-Seq have shown that G4s are preferentially found at gene promoters of transcriptionally active genes, acting as transcriptional enhancers rather than repressors (Di Antonio et al., 2020; Hansel-Hertsch et al., 2016; Hansel-Hertsch et al., 2020; Lago et al., 2021; Zheng et al., 2020). Moreover, G4 formation seems to contribute to activated gene expression by means of many different mechanisms including regulating chromatin architecture, stabilizing R-loops, promoting long-range DNA interactions, triggering liquid-liquid phase separation, direct transcription factor binding, and via promoting guanine oxidation and repair (Robinson et al., 2021).

In the latter cases, including the transcriptional enhancement observed in response to oxidative damage of the G4 *VEGF* promoter, BER proteins and their repair activities are a key part of the mechanism of G4-associated gene regulation. Because of the key role played by BER proteins in reading the damage as an epigenetic mark, recruiting the gene-activating transcription factors, and ultimately repairing the damage, we sought out to further our understanding of how this is damage is repaired. Furthermore, beyond gaining insight into the mechanism of transcription regulation, understanding how BER in the context of G4s (which are oxidative damage hotspots) is relevant as the process is key for avoiding accumulation of the mutagenic lesion, 8oxoG, which can lead to G to T mutations via mispairing with adenine. Interpreting the results of our BER assays reported here in the context of the published literature, we propose a model for BER in the context of the *VEGF* G4 promoter, Figure 7.

**Figure 7:**
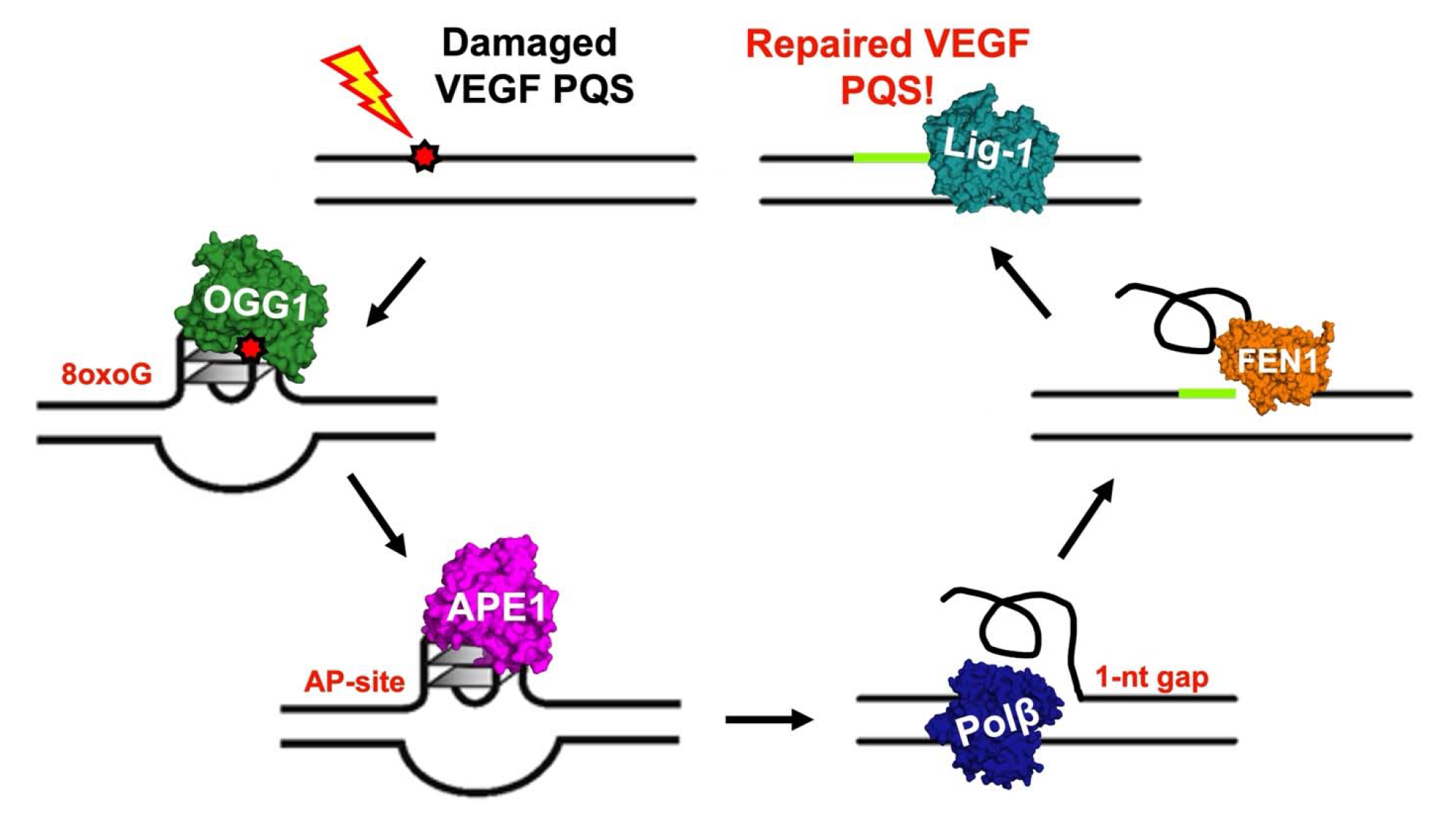
Model for BER of the *VEGF* G4 promoter DNA sequence.

The *VEGF* promoter PQS is susceptible to oxidative damage in both its B-form and G4 conformations (Fleming et al., 2015) and it remains unclear at what stage of the 8oxoG induced, BER-dependent gene enhancement mechanism the DNA adopts the required G4 fold. Prior studies utilizing single-stranded DNA G4 substrates have consistently revealed a lack of and/or very weak OGG1 glycosylase activity toward G4 DNA containing 8oxoG (Ferino & Xodo, 2021; Zhou et al., 2015). Moreover, OGG1 has been shown not to have activity on non-G4 single-stranded DNA substrates as well (Franck et al., 2022). In addition, based on changes in FRET signals during stopped flow kinetic analysis it has been asserted that while OGG1 can bind single-stranded G4 DNA, that it cannot form the appropriate contacts to induce 8oxoG eversion (Kuznetsova et al., 2020). Combined, this has led to the assumption that the G4 fold is adopted by the *VEGF* PQS after OGG1 activity. Here, we examined OGG1 activity on a double-stranded *VEGF* G4 promoter substrate, which mimics the genomics state with both DNA strands, and show that compared to the corresponding single-stranded substrate, there is much greater product formation (Figure 2). Therefore, the model presented here includes the ability of OGG1 to process lesions in a folded G4 substate. However, additional experiments capable of monitoring G4 folding during OGG1 catalysis are required to determine whether OGG1 cleaves the substrate by first resolving the G4 structure, or if the presence of the opposing strand or flanking duplex regions allow for its enhanced activity on our double-stranded substrates. Of note, Pol β DNA synthesis of the double-stranded *VEGF* G4 substate generated by OGG1/APE1 has the same G4-substrate specific multiple insertion pattern as the Pol β substrates originating as G4s with 1-nt gaps, Figures 4 and 5. Therefore, If OGG1 does resolve the G4 it is likely re-folded during a subsequent step of BER (i.e., by APE1, see below).

APE1 can bind and stabilize G4 DNA (Fleming et al., 2021; Roychoudhury et al., 2020). Therefore, if OGG1 does bind and subsequently resolve the G4 before cleaving the N-glycosyl bond, APE1 likely re-folds the G4 structure while bound. Moreover, prior work has shown that APE1 activity is greatly inhibited by G4 DNA despite tight binding affinities for the structure (Fleming et al., 2021; Sowers et al., 2023). Because pre-steady-state kinetics enable the resolution of distinct key catalytic steps (i.e., DNA cleavage and product release) and we utilized double-stranded AP-G4 *VEGF* promoter substrates, the work here expands on these prior observations. Unexpectedly, we observed APE1 to have a 2-fold faster rate of cleavage (k_obs_) on double-stranded G4 DNA *VEGF* substrates compared to B-form DNA. On the other hand, more in line with the steady-state assays published previously, we do see a decrease in the steady-state rate (k_ss_) of APE1 on double-stranded *VEGF* G4 DNA substrates, although the degree of the decrease is far less than shown in the past on single-stranded *VEGF* G4 DNA substrates. Importantly, like others, we also observed the greatly reduced APE1 activity and product yield in assays utilizing single-stranded *VEGF* G4 DNA substates (Figure 5, lane #2 in the second vs. third time courses). This new insight on APE1 catalytic activity, including the higher than anticipated rates of DNA cleavage and reduced product release on the double-stranded *VEGF* G4 promoter DNA substrate, is particularly relevant toward understanding the role played by APE1 in regulating *VEGF* transcriptional regulation, as its slow repair activity on the G4 DNA has been proposed as a key part of the mechanism for transcriptional activation. Specifically, the APE1-G4 binary complex has been proposed to serve as a hub for the recruitment of activating transcription factors to induce mRNA synthesis at least in part due to its reported slow repair activity on the G4 DNA substrates (Fleming et al., 2017; Howpay Manage et al., 2023). However, considering these new findings, perhaps the role played by APE1 is more complex and other protein factors and/or APE1 PTMs are required to switch APE1 out of repair and into a transcription modulator. For example, cystine oxidation is considered one of the factors that may trigger for this switch, as cystine oxidation to sulfenic acid has been shown to enhance APE1 affinity for G4 DNA while broadly reducing its repair activity (Howpay Manage et al., 2022), however this effect has yet to be studied in the context of double-stranded G4 substrates. Along similar lines, acetylation of APE1 in chromatin has been demonstrated to delay its dissociation from the AP site (Roychoudhury et al., 2020).

After APE1 strand cleavage, Pol β is tasked with removing the cleaved AP site with its AP lyase activity and gap filling DNA synthesis. To our knowledge there are not any published studies looking at DNA Pol β activity in the context of G4 DNA. However, the impact of G4 DNA sequences and topology on the fidelity of other human DNA polymerases has been studied. These works have demonstrated unique polymerase error signatures during the synthesis of G4 motifs within, immediately flanking, and encompassing the G4 motif depended on the particular G4 topology and DNA polymerase (Stein et al., 2022). Like these studies, we saw changes in Pol β activity with the *VEGF* G4 DNA substrates compared to B-form DNA, with Pol β inserting more than one nucleotide via strand displacement DNA synthesis in the case of the *VEGF* G4 DNA substrates. We attribute the multiple insertions by Pol β to strand displacement DNA synthesis indicative of long-patch BER for several reasons. First, despite the DNA adopting a G4 structure, several of our results indicate that Pol β utilizes the DNA template. This includes preferential insertion of dGTP when C is in the template position, and the requirement for a double-stranded substrate. Second, for the PQS to adopt a G4 structure, as our CD data indicates it does, the strand must become displaced from the template. This is similar to other conditions that lead to strand displacement DNA synthesis, such as in substrates with a deoxyribose resistant to β-elimination (Podlutsky et al., 2001). Moreover, strand displacement DNA synthesis is also seen in trinucleotide repeat (TNR) sequences, which undergo spontaneous strand slippage resulting in larger gaps and the formation of non B-form DNA structures including DNA hairpins (Liu et al., 2009). Since DNA synthesis occurs via extension of the 3’ end of the DNA, this would indicate that the 3’ end of the DNA preceding the site of damage in the 1-nt gapped substrate is able to re-anneal to the DNA template strand. It remains unclear how and when this occurs, but the downstream strand could remain in a non B-form structure providing the larger gap and promoting the long-patch sub-pathway of BER. It also remains unclear whether DNA damages located more 3’ within the G4 DNA than positions 7 and 12 would represent less efficient substates for Pol β due to either challenges related to the upstream strand re-annealing to the template or a lack of non B-form secondary structure to promote strand displacement in the downstream strand.

Additional support for the long-patch BER model is the observed requirement for FEN1 in the final step of BER, DNA ligation (Figure 6). Of the two damage positions, FEN1 allowed for more ligation in the substrate with damage at position 12 than that with damage at position 7. This may be the result of the shorter flap generated with the substate, due to the shorter G-run associated with position 7 and limiting the reaction to dGTP insertion. If this is the case, since all four nucleotides would be present in the biological scenario, strand displacement DNA synthesis would potentially generate longer flaps in the case of both substrates, which would potentially favor FEN1 activity and allow for more efficient ligation. In fact, we do see increased insertions when we give the reaction all four nucleotides (Supplemental Figure 3).

While the requirements for OGG1 and APE1 in the transcriptional regulation of *VEGF* have been clearly established (Fleming et al., 2017; Fleming et al., 2021), it remains unclear if the downstream factors also play a role in the transcriptional regulation. With the observation that APE1 cleaves and releases G4 products with close to same efficiency as it does B-form DNA a possible role for the other factors becomes more plausible, as the mode of switching between the two activates may be more complicated than prior thought. For example, in cases where DNA synthesis and/or ligation are confounded by the G4 substrates relative to B-form substrates, the nicked G4 DNA could result in additional binding of APE1 during which it could potentially recruit activating transcription factors.

### Statement of Author Contributions

Dr. Amy Whitaker, Adil Hussen, and Haley Kravitz analyzed the data and prepared draft figures and tables. Dr. Amy Whitaker prepared the manuscript draft with intellectual input from Dr. Freudenthal. All authors approved the final manuscript.

## Acknowledgements

We thank Dr. Bill Beard for sending us the plasmid for DNA ligase 1, FEN1 protein and important intellectual input.

