## Supplemental for "The *VEGF* G-quadruplex forming promoter is repaired via long-patch BER"

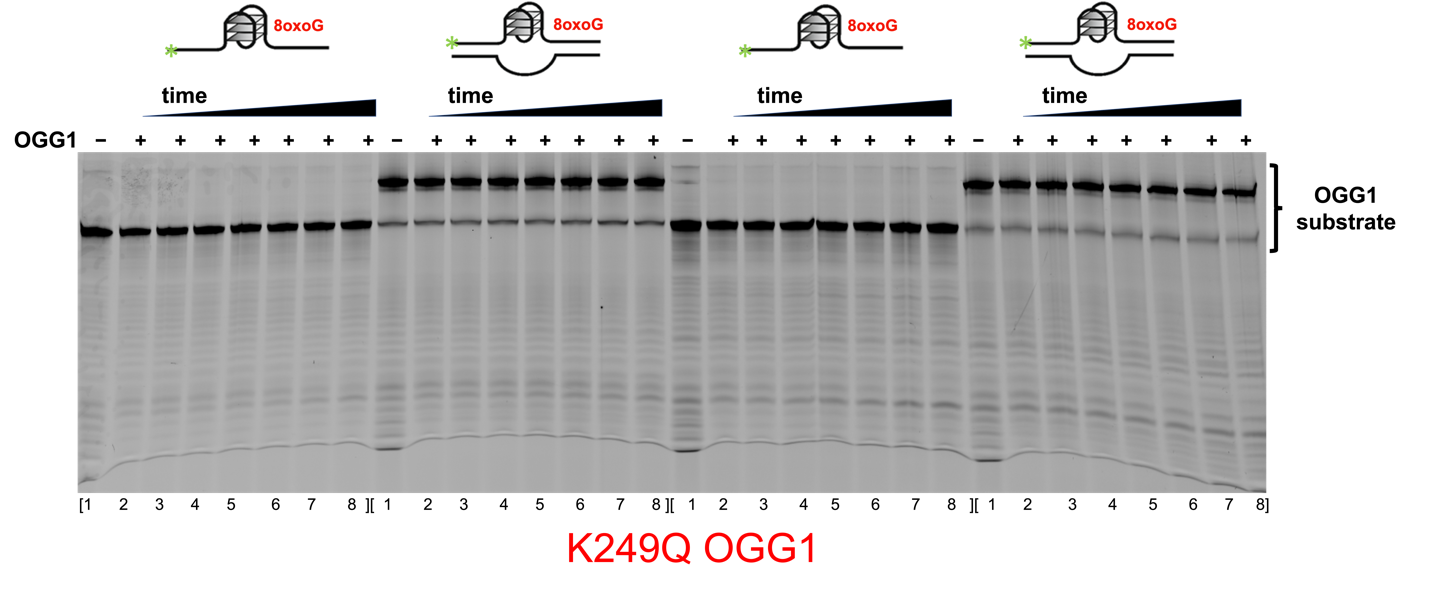

**Supplemental Figure 1:** K249Q OGG1 cleavage of *VEGF* G4 promoter substates. OGG1 reaction time courses on various G4 substrates listed from left to right: the double-stranded *VEGF* G4 with an 8oxoG lesion at position 7, single-stranded *VEGF* G4 with an 8oxoG lesion at position 7, double-stranded *VEGF* G4 with an 8oxoG lesion at position 12, single-stranded *VEGF* G4 with an 8oxoG lesion at position 12. Lane 1 of each panel is a no enzyme control. Lanes 2 – 8 are the reaction products of OGG1 N-glycosyl bond cleavage followed by phosphodiester backbone incision over a 60-minute time course. The reaction time points corresponding to lanes 2 – 8 are 0.5, 1, 2.5, 5, 10, 30, and 60 minutes. Note that the 8oxoG containing double-stranded substrates often run as two species due to partial re-annealing of the complementary strand to the 6-FAM labeled G4 containing strand during gel loading.

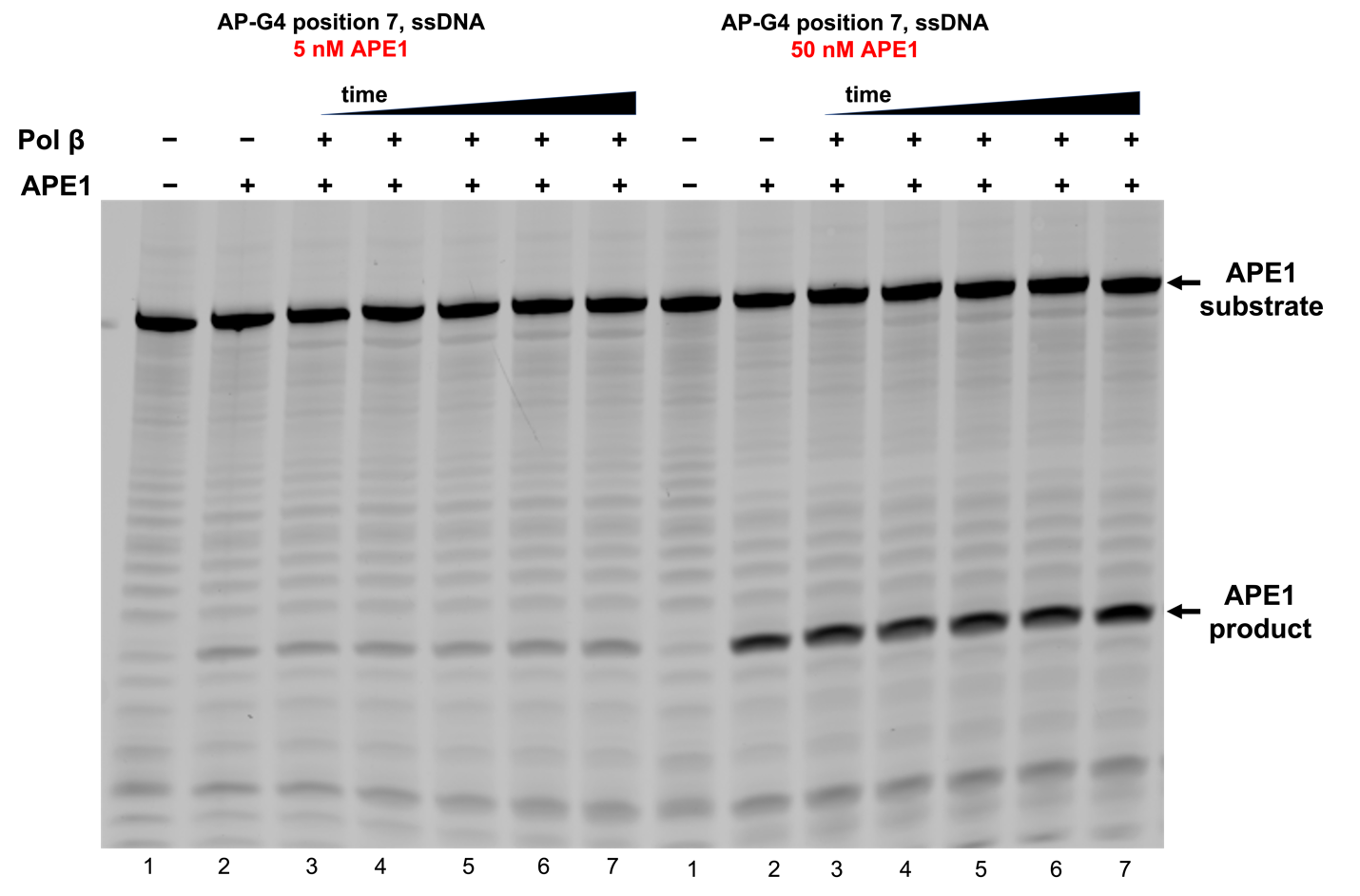

**Supplemental Figure 2:** Representative gel image for Pol β reaction time courses on a single-stranded *VEGF* G4 promoter substrate with a THF at position 12. The substrate was first incubated with APE1 for 60 minutes to generate a substrate for Pol β. Lane 1 of each time course is a no enzyme control. Lane 2 is after 60-minute incubation with either APE1 (5 nM left and 50 nM right). Lanes 3 – 7 are the reaction products of Pol β insertion of dGTP over a 60-minute time course. The reaction time points corresponding to lanes 3 – 7 are 2.5, 5, 10, 30, and 60 minutes.

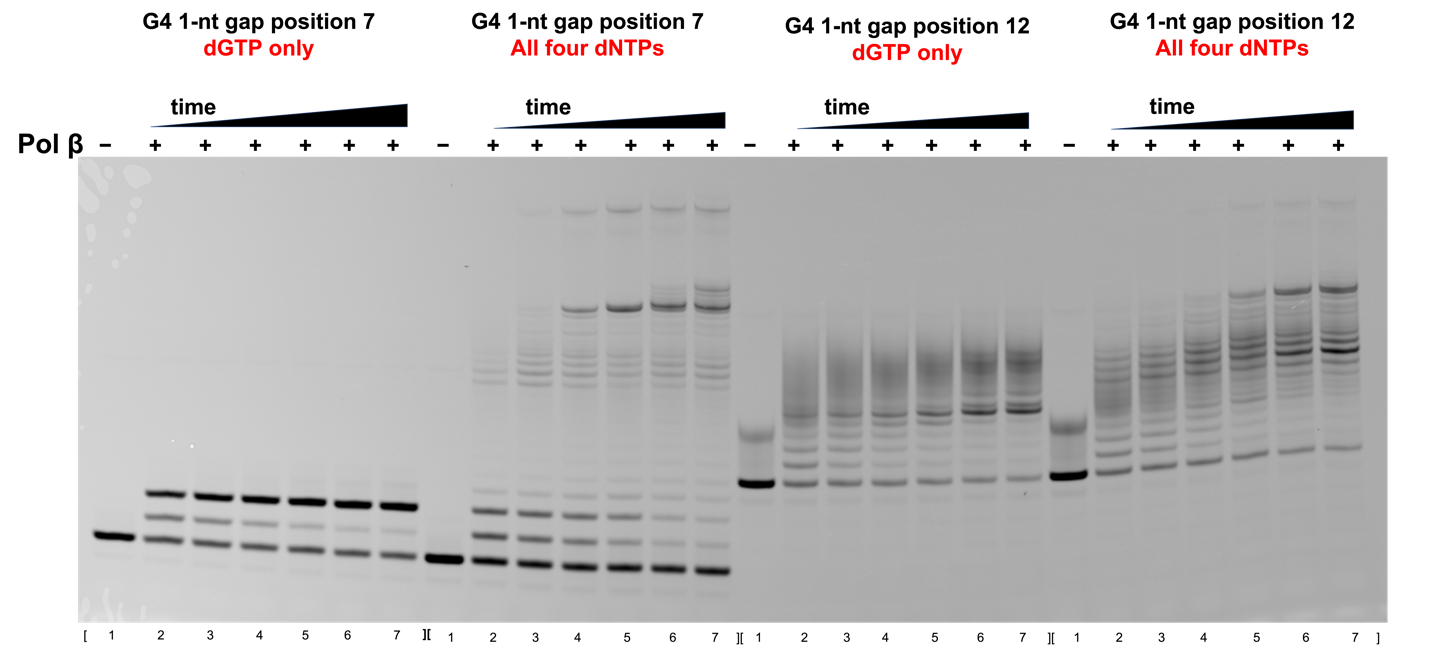

**Supplemental Figure 3:** Representative gel image for Pol β reaction time courses on double-stranded *VEGF* G4 promoter substrates a 1-nt gaps at either position 7 or 12. Lane 1 of each time course is a no enzyme control. Lanes 2 – 7 are the reaction products of Pol β insertion of the indicated dNTP (either only dGTP or a pool of dGTP, dATP, dCTP and dTTP) over a 60-minute time course. The reaction time points corresponding to lanes 2 – 7 are 1, 2.5, 5, 10, 30, and 60 minutes.
